# A Pseudo-Temporal Causality Approach to Identifying miRNA-mRNA Interactions During Biological Processes

**DOI:** 10.1101/2020.07.07.192724

**Authors:** Andres M. Cifuentes-Bernal, Vu VH Pham, Xiaomei Li, Lin Liu, Jiuyong Li, Thuc Duy Le

## Abstract

**Motivation:** microRNAs (miRNAs) are important gene regulators and they are involved in many biological processes, including cancer progression. Therefore, correctly identifying miRNA-mRNA interactions is a crucial task. To this end, a huge number of computational methods has been developed, but they mainly use the data at one snapshot and ignore the dynamics of a biological process. The recent development of single cell data and the booming of the exploration of cell trajectories using “pseudo-time” concept have inspired us to develop a pseudo-time based method to infer the miRNA-mRNA relationships characterising a biological process by taking into account the temporal aspect of the process.

**Results:** We have developed a novel approach, called pseudo-time causality (PTC), to find the causal relationships between miRNAs and mRNAs during a biological process. We have applied the proposed method to both single cell and bulk sequencing datasets for Epithelia to Mesenchymal Transition (EMT), a key process in cancer metastasis. The evaluation results show that our method significantly outperforms existing methods in finding miRNA-mRNA interactions in both single cell and bulk data. The results suggest that utilising the pseudo-temporal information from the data helps reveal the gene regulation in a biological process much better than using the static information.

**Availability:** R scripts and datasets can be found at https://github.com/AndresMCB/PTC

## 1 Introduction

Identifying the regulatory relationships between microRNAs (miRNAs) and messenger RNAs (mRNAs) characterising a biological process is a crucial task. For example, Gregory et al.[1] have found that miR200c regulates ZEB1 during Epithelial to Messenchymal Transition (EMT), a key process in cancer metastasis, and therefore miR200c and its interaction with ZEB1 can be used for the treatment for cancer metastasis. Understanding the miRNA-mRNA interactions that characterise different biological processes will give insights into the biology of diseases and help with the design of treatments of the diseases.

There has been a huge number of computational methods developed to find miRNA-mRNA relationships from data. The first wave of methods looked into the negative correlation in expression levels of miRNAs and mRNAs, as the majority of miRNAs down-regulate their target mRNAs (see [2] for a review). The second wave of methods argued for the causal relationships between miRNAs and mRNAs and aimed to exclude the spurious relationships between miRNAs and their target mRNAs found from data [3,4]. The third and current wave of research is focused on the relationships between miRNAs and mRNAs in relation to other types of molecules, e.g. long non-coding RNAs. An example of this category is the miRNA sponge interaction networks for which researchers investigate the competition between different RNAs, including mRNAs, lncRNAs, and pseudo-genes, in fighting for a place for binding with miRNAs [5, 6].

Most of the current computational methods for miRNA-mRNA interaction discovery use the data generated at one specific snapshot of a biological process and ignore the dynamics or the temporal aspect of biological processes. For example, several works [7, 8] aim to find the miRNA-mRNA interactions by analyzing gene expression data obtained from cancerous tumors. In these works, data employed was collected from different patients at one single time point during their disease. Other works [9, 10, 11] use genetic information, like gene expression, in two different time points or conditions, in order to contrast the findings at different biological states, e.g. cancer vs normal; Epithelial vs Mesenchymal.

However, ideally, we would like to understand what is happening *during* a biological process, e.g. how genes interact with each other during the process when cells transform from normal stage to invasive stage. Time series data collected during a biological process can be used to infer the miRNA-mRNA interactions characterising the process [12, 13], but this dynamic analysis is rarely done in practice. Given a biological process of interest, one needs to collect expression data for both miRNAs and mRNAs at different time points of the process. Due to the costs of the experiments, there are very few such datasets available. Therefore, we have to resort to other types of data and develop novel methods to examine the miRNA-mRNA interactions during a biological process.

Recent development in single cell sequencing and the concept of *pseudo-time* have opened the door for developing novel methods for revealing what is happening during biological processes [14]. Single cell sequencing technique quantifies gene expression profiles of transcripts in individual cells. Meanwhile, the pseudo-time approach utilises the expression of the markers of a biological process to order the cells along the progress of the biological process.

Inspired by the pseudo-time concept [14], in this paper we develop a novel approach, called the *pseudo-time causality (PTC)* based approach, to elucidate the miRNA-mRNA interactions during biological processes, using gene expression data with the expression profiles of matched miRNAs and mRNAs in the same cells or samples. Given a biological process, *PTC* firstly transforms the matched miRNA and mRNA single cell gene expression data to pseudo-time data using the marker genes of the biological process. *PTC* relies on the causal invariance property [15, 16] to find the causal relationships between miRNAs and mRNAs from the pseudo-time data. We have applied *PTC* to the single cell dataset from [17] and the bulk data from [18] (please refer to sections 3.1 and 3.5 for a detailed description). In both cases, VIM, an EMT marker, is used to define the pseudo-time.

The results have shown that *PTC* significantly outperforms the benchmark methods in identifying experimentally confirmed miRNA-mRNA interactions using either single cell or bulk data. The results suggest that the temporal information during a biological process is useful for revealing the miRNA-mRNA interactions characterising the biological process.

## 2 Methods

### 2.1 Problem definition and method outline

Based on the premise that a biological process can be characterised by miRNA-mRNA interactions, it becomes necessary to develop a method capable of inferring those interactions from data obtained during the biological process, ideally from time-series data obtained from experiments carried out during the biological process of interest. Unfortunately, collecting time-series experimental data is costly and sometimes unfeasible. Because of that, most of the current experimental data is static data obtained in few times points during a biological process.

*PTC* relies on the assumption that a biological process, like cancer progression, is governed by an underlying causal mechanism throughout. This presumption defines the two main research problems to be solved in this paper. Firstly, temporal data from different time points during the process is required in order to infer the miRNA-mRNA relationships characterising the process. Secondly, since miRNA-mRNA relationships are considered causal, a way to test causality during the biological process is also required.

*PTC* solves the temporal data requirement by performing a pseudotime analysis and transforming static data (in our case, miRNA/mRNA gene expression) to their pseudotime ordered version. In order to determine the causal relationships between miRNAs and mRNAs, our method presumes that the relationship between a mRNA (the target) and a set of miRNAs (the regulators) can be modeled as a linear system where the effect or response variable corresponds to the gene expression of the mRNA and the cause or predictor variables correspond to the expressions set of miRNAs.

As shown in Figure 1, the overall procedure of our proposed method, *PTC*, consists of three phases:

- **Phase 1: Pseudotime analysis.** *PTC* takes a (static) gene expression dataset containing samples of matched miRNAs and mRNAs gene expression and converts the dataset into a time series dataset. The conversion is done through a pseudotime analysis to reorder the samples according to the pseudotime established using a well recognised EMT marker, VIM (details are in Section 2.3).
- **Phase 2: Gene selection**. This phase consists of two steps, a pre-selection process and a process to define the set of plausible predictors or causes for a mRNA. In the first step, the top ranked miRNAs and mRNAs based on their gene expression median absolute deviation (*MAD*) are selected. In the second step, for each of the selected mRNAs, a further analysis is carried out to determine those miRNAs that can biologically interact with the mRNA, and only the miRNAs that can target the mRNA as predicted by TargetScan [19] are kept as the plausible predictors or causes of the mRNA.
- **Phase 3: Identification of miRNA-mRNA**. In this phase, *PTC* employs the *decoupled test* [16] to determine whether the set of plausible predictor miRNAs of a mRNA obtained in Phase 2 are the real causes or regulators of the mRNA or not. To determine the causal relationships, *PTC* tests if the “causal invariance” property is violated by the relationships during the biological process by using the time series data obtained above with the gene selection in Phase 2 (Details are in Section 2.4).

**Figure 1:**
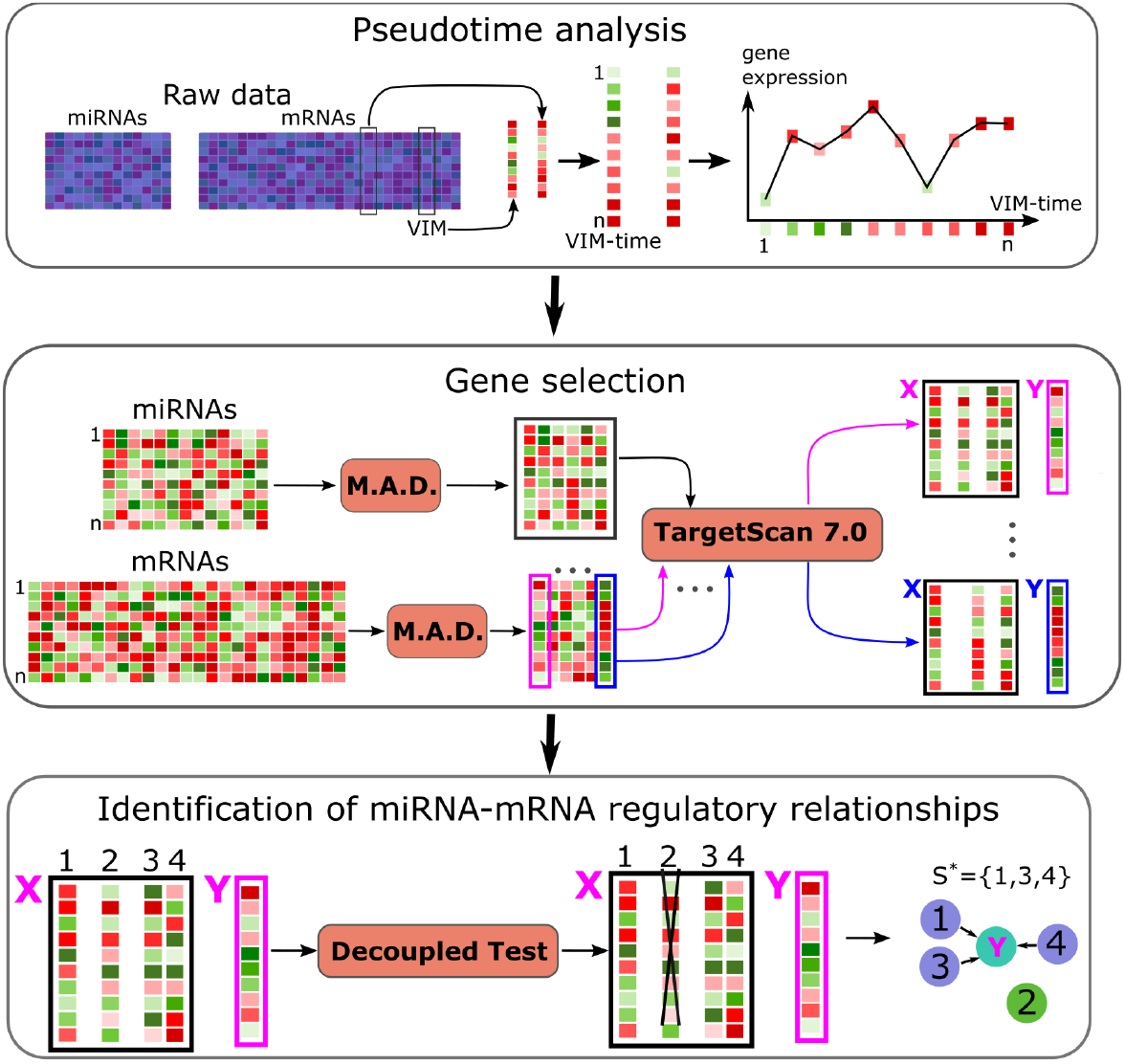
Summary of the developed method, *PTC (Pseudo-Time Causality).* PTC comprises three phases: (1) Pseudotime analysis. miRNA-mRNA static gene expression is processed to create time series data following a pseudotime order. In our experiments, VIM was selected as the biomarker to create the pseudotime. (2): Gene selection. MAD and TargetScan 7.0 (conserved sites predictions) are employed to determine a set of miRNAs as plausible predictors of each of the **k** genes. (3) Identification of miRNA-mRNA regulatory relationships. Violations to the invariance property are assessed to find the causal parents (predictors) of each gene.

### 2.2 Pseudotime Analysis

Since most gene expression datasets contain only a snapshot of the expression of genes, it becomes necessary to make a data transformation for temporal analysis. Our method relies on the assumption that dynamic causation can be inferred from the underlying dynamic in temporal data. In general, it is desired to find those mRNAs whose gene expression changes progressively with time, and use these genes as pseudo temporal units to model the different trajectories during cancer progression of all genes in the genomic data.

In our experiments, the biological process selected is the EMT process. We use VIM to define the pseudotime as VIM is a well-known EMT marker (Puchinskaya, 2019; Shu et al., 2019). During epithelial-mesenchymal transition (EMT), the gene expression level of VIM increases [20]. This implies that the ascending ordering of the expression of VIM in an EMT dataset provides us a pseudotime of EMT, called VIM-Time in this paper. Hence to convert the static matched mRNA and miRNA samples into temporal data, in our method we sort the matched mRNA and miRNA samples based on the VIM-Time, i.e. we sort the input data matrix in ascending order of the column containing VIM’s expression values. As a result, changes of gene expression of all genes were described as a progression in terms of VIM-time. This data transformation incorporates a temporal dimension to the data that *PTC* uses to identify causal relationships during the EMT process.

### 2.3 Gene selection

A comprehensive analysis to determine all parents of a response variable by testing the invariance property, could require evaluating all possible combinations of the plausible predictors. Peters et al. proposed to determine the set of true causes of a variable as the intersection of all sets that do not violate the invariance property [15]. Finding the intersection set could imply to test a huge number of combinations. This can easily be unfeasible if the number of plausible predictors is big. For example, if an mRNA has 30 plausible predictors, in Phase 3, in the worst case, for this single mRNA we will need conduct over a billion tests since there are 2^30^ possible combinations for 30 miRNAs.

Taking this fact into consideration, in our method the gene selection phase includes a heuristic procedure to reduce the number of predictors for each gene. First, a subset of miRNAs and a subset of mRNAs are selected during the first step of this phase. This selection is based on the hypothesis that miRNAs and mRNAs with most expression variability are more likely related with regulation processes in cancer. For **PTC**, we choose to use *Median Absolute Deviation (MAD)* as the measure for gene expression variability. This selection was performed using the function *“FSbyMAD”* from the *“CancerSubtypes”* R package [21]. In our experiments, parameters for *“FSbyMAD”* were: cut.type = “topk”, value = 30 when selecting miRNA and 1500 when selecting mRNA, replicating the selection process described in [18].

After the above selection step based on *MAD, PTC* employs a heuristic to further reduce the set of plausible predictors for each mRNA, to the set of miRNAs that can bind that gene as predicted by TargetScan 7.0. The set of plausible predictors of each gene is restricted to a tractable size for the testing in Phase 3.

### 2.4 Identification of miRNA-mRNA regulatory relationships

It has been shown that causal relationships have an “invariance” property, that is, for a causal structure, the conditional distribution of a variable (called a target or effect variable in this case) given all its direct causes remains the same under interventions on any other variables in the structure except the response variable. This “property” has been discussed in literature under different names like “autonomy” “stability” or “modularity” [22, 23, 24, 25, 15].

This property of invariance can be exploited to determine whether a set of plausible predictors is the set of direct causes of a response variable. In general terms, given a set of plausible predictors and one response variable, violations to the invariance property can be assessed using data obtained from different environments (which may be associated with interventions on different variables as described above). If a set of plausible predictors violates the invariance property across different environments, it can be inferred that the set of plausible predictors contains variables which are not direct causes of the response variable.

#### 2.4.1 Testing violation to the invariance property

In phase 3, our method determines the existence of a causal link between a set of miRNAs (parents) and a mRNA by assessing violations to the invariance, adapting the definition of “invariant set” given in [16] as follows:

#### Definition 2.1

***(Invariant set)*.** Given a column vector **Y**≔ ((*Y_t_*)^*T*^)*t*∈{1,..,*n*}, *containing the gene expression for n temporally ordered samples of the mRNA considered as the response variable of interest, and a matrix* 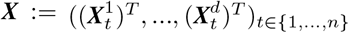, *containing the gene expression for n temporally ordered samples of the d miRNAs considered as plausible predictors of the mRNA, after the phase 2 of PTC. Furthermore, considering that for any set S* ⊆ {1,..., *d*}, *the matrix **X**^S^ contains only the columns of **X** indexed by S. S is an “‘invariant set” with respect to* (***Y***, ***X***) *if there are parameters μ* ∈ ℝ, *β* ∈ (ℝ \ {0})^|*S*|×1^ *and σ* ∈ ℝ_>0_ *such that*:

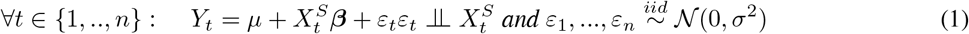

*Where β is the vector of regression coefficients, μ is the regression independent term, and ε*_1_…, *ε_n_ are independent and identically distributed random noises, following a normal distribution with mean 0 and standard deviation σ.*

Based on the above definition of an invariant set, if S does not violate the invariance property, ∀*t* ∈ {1,.., *n*}, *μ, β* and *σ*^2^ remain the same. This feature allows the use of m adjacent time points, with *m* < *n*, to create segments time series and use them as environments, instead of data from different experimental settings. It is important to note that the segments of the time series that simulate a single environment must have the same number of time points, and they must be synchronized.

*PTC* searches for the following violations in the models of Y (as in Eqn. 1) inferred from different segments of the time series: (i) notable differences between the vectors of regression coefficients (*β*) and/or (ii) differences in the noise variance (*σ*^2^). For experiments in Section 3, we divide the time series in two similar long segments to compare and assess differences in the invariant set models.

The *decoupled test* [16], so named because it test the two violations described above independently, is employed to verify if a set of miRNAs commit any of the violations to the invariance described above. To conduct the test we use the function *seqICP.s* from the seqICP R package [16], with a significance level *α* = 0.02. The output of this test is the lowest of the p-values obtained after testing the two violations described above. In our experiments, a p-value < 0.02 implies at least one violation to the invariance property.

#### 2.4.2 Identifying miRNAs regulating a mRNA

For each mRNA of interest (Y) and its set of plausible predictor miRNAs obtained during Phase 2, *PTC* conducts the above described invariance property test on each subset of the set of plausible predictors to find all the subsets of miRNAs that do not violate the invariance property. Then *PTC* outputs the union of all the subsets which possess the invariance property (denoted as *S*\*), as the set of direct causes or regulators of Y. The decision of using the union of all subsets of plausible predictors which do not violate the invariance property is based on the fact that *PTC* only admits as plausible predictors those miRNAs that can biologically target the gene.

To make *PTC* more efficient, the test of invariance property starts with the testing on the largest subset of the d plausible predictors, i.e. {1,.., *d*}, then we test the subsets of size (d-1) and so on, and finally the subsets of size 1. If *S*′ ⊆ {1,2,..., *d*} is tested to satisfy the invariance property, then all subsets of *S*’ are not tested, since *PTC* outputs the union of all subsets of {1,2,..., *d*} satisfying the invariance property. For example, assume that a gene has three plausible predictors, {1,2,3}. In this instance, the tests would be conducted in the following order: {1,2,3} {1,2} {1,3} {2,3} {1} {2} {3}. If {1,3} is tested not to violate the invariance property, then all of its subsets, i.e. {1} and {3} will not be tested.

The final output of *PTC* contains all the identified miRNA-mRNA pairs and their ranking scores. We use the TargetScan 7.0 Context++ scores for all conserved miRNA sites and the score of each miRNA-mRNA pair was calculated as the average of all context scores of that pair across all conserved sites. Since context scores are negative numbers, the more negative the average is, the higher the rank the miRNA-mRNA pair has in our method.

The performance of *PTC* is measured by the percentage of miRNA-mRNA relationships inferred by *PTC* that have been confirmed experimentally. For the evaluation, a collection of confirmed interactions has been assembled by combining information from several databases, including miR-Tarbase 6.1 [26], Tarbase 7.0 [27] and miRWalk 2.0 [28]. An inferred interaction is considered as experimentally confirmed if it can be found in at least one of the above databases. The performance of *PTC* is also compared against traditional miRNA-mRNA interaction inference methods, including *Pearson* [29] and *Lasso* [30], by checking the proportion of confirmed interactions predicted by them. “GO biological processes” and “KEGG pathways” analyses were performed on the miRNA-mRNA causal relationships determined by *PTC* to assess their relevance to cancer and EMT.

## 3 Results and discussion

### 3.1 Identified miRNA-mRNA interactions in EMT with single cell data

The technique of quantifying the expression of miRNAs and mRNAs in single cells at the same time is still in its infancy, as single cell techniques normally destroy the cell when measuring the gene expression and thus is hard to measure the miRNA expression at the same time. Currently, only one work [17] provides matched miRNA and mRNA expression for single cells, but with very limited number of cells (20 cells). Gene Expression in this data set comes from single human acute myeloid leukemia cells.

We apply *PTC* to this single cell dataset [17] to find miRNA-mRNA interactions in EMT. This dataset provides a total of 20 cells that were each split into halves in order to obtain paired miRNA - mRNA gene expression data (miRNA expression from one half cell and mRNA expression from the another half of the same cell). In this research only 19 cells that have all paired entries were employed, with cell number 6 excluded from the analysis since it does not have paired miRNA-mRNA gene expression. After processing (please see Supplementary Material, Section 1), our data set has 2822 miRNAs and 23141 mRNAs from 19 single cells.

From the above processed dataset, *PTC* has predicted 569 miRNA-mRNA interactions as visualised in Figure 2. Experimentally verified interactions are shown in red. All other inferred regulatory relationships are represented in blue, and their strength is indicated by color intensity. The darker the link the stronger the predicted interaction is.

**Figure 2:**
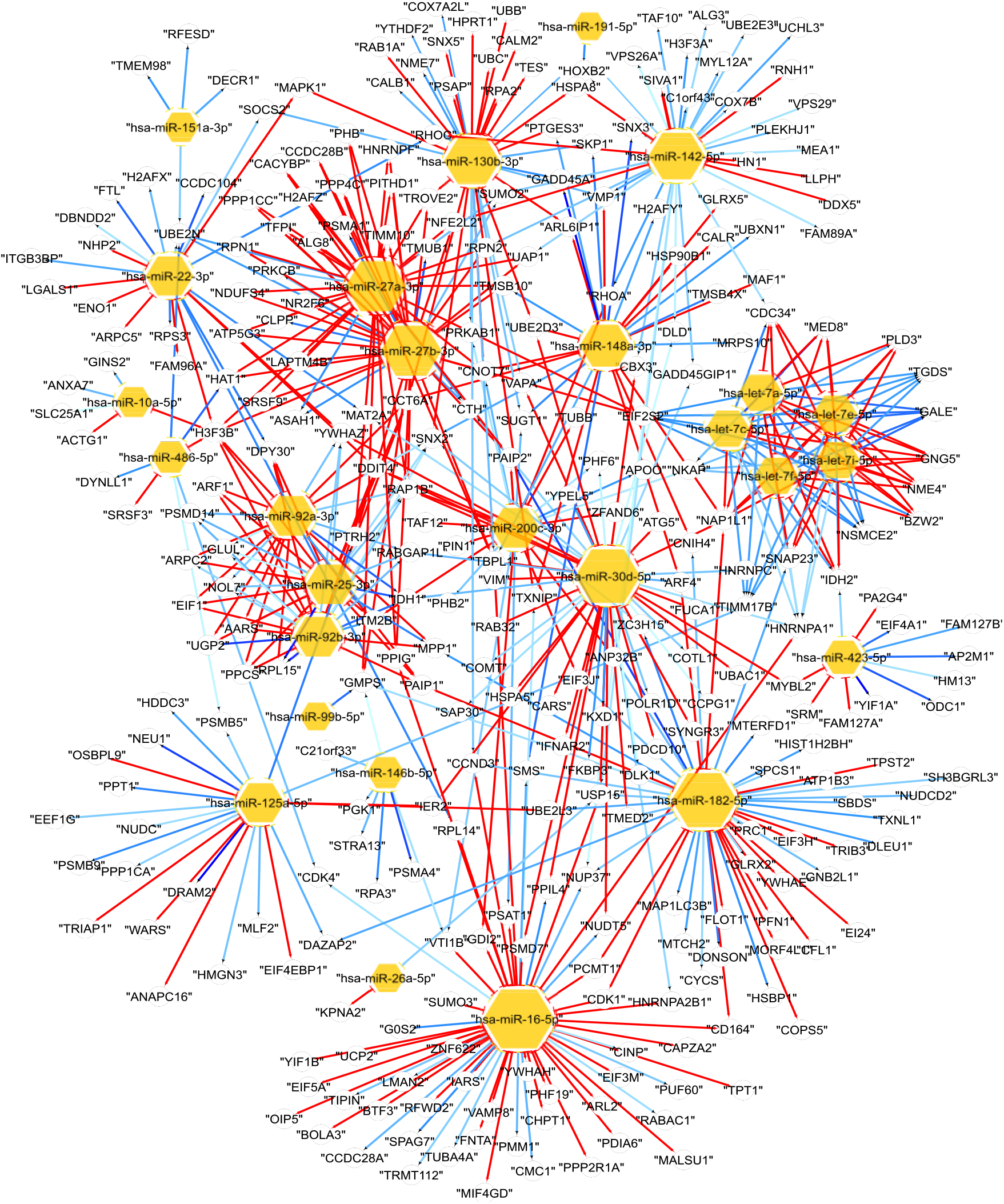
Regulatory relationships discovered by PTC. In red, experimentally confirmed interactions. In blue, all other predicted interactions. Strength of the causal link is represented by color intensity, with darkest color representing the stronger links.

The number of genes predicted to be regulated by a miRNA is indicated by the size of the miRNA node, i.e. a larger sized miRNA node has more predicted targets. “hsa-miR-16-5p”, “hsa-miR-182-5p”, “hsa-miR-30d-5p”, “hsa-miR-27a-3p” and “hsa-miR-27b-3p” were identified as the most influential nodes, with 55, 47, 46, 40 and 40 inferred regulatory relationships, respectively. Several studies have linked these miRNAs to different types of cancer. “hsa-miR-16-5p” appears to have a significant role in chronic lymphocytic leukemia [31] and periocular Sebaceous Gland Carcinoma [32]. “hsa-miR-182-5p” has been related with colorectal cancer [33], drug resistance in breast cancer cell lines [34], Epithelial Ovarian Cancer [35] and lung adenocarcinoma [36]. “hsa-miR-30d-5p” has been linked with nonmuscle invasive bladder cancer [37], while “hsa-miR-27a-3p” with Spinal Cord Glioma Progression, intrahepatic cholangiocellular carcinoma [38]. Finally, “hsa-miR-27b-3p” has been related to tumor suppression in lung cancer [39] and gastric cancer [40].

### 3.2 A significant number of interactions discovered by *PTC* has been experimentally confirmed

Among the 569 miRNA-mRNA regulatory relationships discovered by **PTC**, 294 pairs are experimentally confirmed interactions. For each group of top interactions predicted (following to the rank explained in Section 2.4.2), the ratio of confirmed interactions increases as follows (Table 1): 121 out of 200 (60.5%), 93 out of 150 (62%), 65 out of 100 (65%), and 37 out of 50 (74%).

**Table 1:** Summary of performance results for *PTC*, Pearson and Lasso. Comparison of confirmed interactions in the top k lists inferred by each method are listed.

| PTC |  |  | Pearson |  | Lasso |  |
| --- | --- | --- | --- | --- | --- | --- |
| Top | Confirmed | Percentage | Confirmed | Percentage | Confirmed | Percentage |
| 200 | 121 | 60.5% | 25 | 12.5% | 28 | 14% |
| 150 | 93 | 62% | 21 | 14% | 21 | 14% |
| 100 | 65 | 65% | 16 | 16% | 14 | 14% |
| 50 | 37 | 74% | 10 | 20% | 8 | 16% |

For comparison purposes, this data set was analyzed with *Pearson* and *Lasso* methods from the miRLAB R package [41]. These two methods are commonly used for inferring regulatory relationships. Inferred interactions by this two methods were verified using the databases of confirmed interactions described previously. The performance of *PTC* is significantly better than the *Pearson* and *Lasso* methods (see Table 1).

### 3.3 *PTC* identifies interactions relevant to EMT

Since *PTC* is based on the idea that there are miRNA-mRNA regulatory relationships that characterize a biological process (EMT process in our experiments), a search was performed to detect relevant interactions during EMT that were inferred by our method. In order to determine if a miRNA-mRNA discovered by *PTC* is EMT related, we assembled a list of EMT miRNAs/mRNAs that have been experimentally confirmed as relevant in the EMT process (please refer to Table S1A. Generic EMT signature for tumour [42], and Table S1. MicroRNAs associated with epithelial-mesenchymal plasticity, [43]). In our experiments a miRNA/mRNA is considered EMT relevant if it can be found in the list explained above.

The numbers of EMT miRNAs in the top 200, 150, 100 and 50 miRNA-mRNA interactions discovered by *PTC* are 12, 11, 10 and 9, respectively. A significant number of the miRNA-mRNA interactions discovered by *PTC* involve these EMT miRNAs. The percentage of relationships inferred by *PTC* that involve those EMT miRNA biomarkers across the tops lists are: 44.5% (89 out of 200), 44% (66 out of 150), 46% (46 out of 100), and 44% (22 out of 50). The inferred pairs containing EMT miRNAs are also highly ranked by our method. It makes sense that the number of EMT miRNAs in inferred pairs by our method is high and persistent because of the fact that our method employs the temporal information during the process of EMT to determine the causal regulatory relationships during the EMT process in cancer.

*PTC* is able to discover EMT related miRNA-mRNA interactions that other methods fail to discover. For example, the experimentally confirmed interaction (hsa-miR-16-5p - VAMP8) was discovered by **PTC**. The classical correlation-based methods *Pearson* and *Lasso* failed to detect this relationship because of its low correlation and “non-significant” p-value. This suggests that, thanks to the fact that *PTC* uses temporal information during the process, it is able to detect causal relationships correlation methods are unable to identify. Figure 3 shows the gene expression of the pair (hsa-miR-16-5p - VAMP8) following the VIM-Time order. An apparent correlation can be observed in some areas of the graph (e.g. VIM-Time between 5 and 10) and some areas show little correlation (e.g. VIM-Time from 10 to 15).

**Figure 3:**
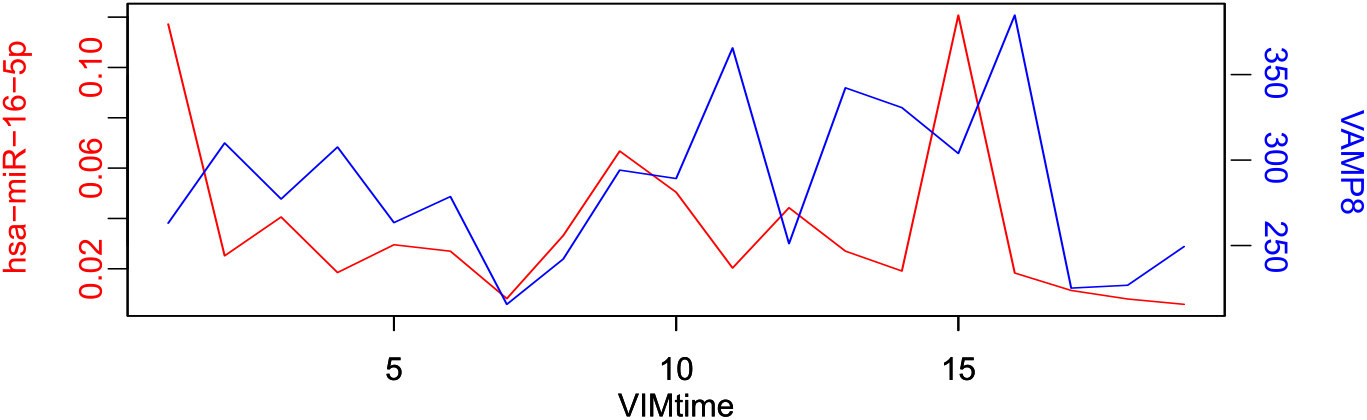
*PTC* can infer EMT miRNA- EMT mRNA that are not discovered by correlation methods. In the figure, gene expression for hsa-miR-16-5p (red) - VAMP8 (blue) following VIMtime order are shown. Correlation between this two expression is 0.04591796 (Pearson’s product-moment correlation, p.value 0.8519262). This regulatory pair is rejected by Pearson but discovered by our method. This regulatory relationship has been experimentally confirmed.

We have also conducted “GO biological processes” and “KEGG pathways” analyses for the miRNAs and genes in interactions discovered by *PTC* to examine their relevance to cancer and EMT. These analyses were performed by using the *Enrichr* web tool [44]. A large number of mRNAs present in the regulatory pairs identified by our method are associated with KEGG pathways and GO biological processes related to EMT process and cancer (please refer to Supplementary Material, Section 2).

### 3.4 *PTC* is stable with different pseudotimes

In order to test the stability of *PTC* with respect to the pseudotime transformations, we also used the *Wanderlust* algorithm [14] to identify EMT pseudotime. *Wanderlust* outcome is a pseudotime based on pre-defined EMT markers from 315 general EMT markers in cancers [42]. In the single cell mRNA data employed in this research, there are 145 of those EMT markers. Thus, we estimate pseudotime by running *Wanderlust* on these 145 EMT markers.

There are four key parameters in *Wanderlust:* the number of nearest neighbors *k*, the shared nearest neighbour *snn*, the number of neighbors selected for each node in a k-nearest neighbors graph *l* and the number of l-out-of-k-nearest neighbors graphs *ng*. In our experiments, the parameter *k* was set to 4 to maintain a reasonable proportion between the number of cells (19) and the number of neighbors per node. Parameters l and ng must be a positive integer lower than *k*. They both were set to 2 since it is the median of the possible options, i.e. {1, 2, 3}. Because all other parameters were fixed, parameter *snn* requires to be set to 0.

The performance of *PTC* when Wanderlust pseudotime was employed is very similar to the *PTC* when VIM-Time was used. 570 miRNA-mRNA relationships were identified when using Wanderlust, in comparison to 569 when using VIM-Time. Confirmed interactions among the top miRNA-mRNA interactions discovered are also similar as shown below: top 200: 122 (Wanderlust) vs 121 (VIM-Time), top 150: 94 (Wanderlust) vs 93 (VIM-Time), top 100: 65 (Wanderlust) vs 65 (VIM-Time), 37 (Wanderlust) vs 37 (VIM-Time).

Only the experimentally confirmed pair (hsa-miR-10a-5p - H3F3B), inferred by *PTC* when VIM-Time was used, could not be detected when *Wanderlust* was employed. Analogously, only the pairs (hsa-miR-27a-3p - ID3), (hsa-miR-27b-3p - ID3), detected by *PTC* when *Wanderlust* was used, were not detected when VIM-Time was employed. The above results suggest that *PTC* is robust and stable to different EMT pseudotime data transformations.

### 3.5 PTC performs better than benchmark methods in bulk gene expression data

Since most methods for inferring gene regulatory relationships use bulk data, and in order to analyze the performance of *PTC* with this kind of data, a benchmark with bulk data was performed. *PTC* was tested using the bulk miRNA and mRNA expression data employed in [18]. This data set contains 518 miRNAs and 17403 mRNAs from 503 tumor samples, downloaded from *“The Cancer Genome Atlas Research Network”* (TCGA) (https://www.cancer.gov/tcga) - BRCA project. The methods selected for comparison were *Pearson* [29], *Lasso* [30], *idaFast* [3], *jointIDA_direct* [45], and the methods based on *Invariant Causal Prediction (ICP) (hiddenICP, hiddenICP pam50, Borda hiddenICP, Borda hiddenICP Pam50),* proposed in [18]. The first four methods were selected because they are widely used in gene regulatory inference. *ICP* based methods were selected because they also use causal invariance property, but with static data.

With this bulk data set a total of 293 miRNA-mRNA regulatory relationships were identified by our method. For all methods, the top 200, 150, 100 and 50 inferred regulatory relationships were selected for analysis. For all the methods but *PTC*, the original research of [18] provides a rank for each interaction inferred by each method, which was used in our analysis for determining the top lists for those 8 methods compared. *PTC* uses it own ranking system, based on the average of the TargetScan 7.0 Context++ scores for all conserved miRNA sites of inferred miRNA-mRNA pairs across all conserved sites (Section 2.4.2). Figure 4 shows the amount of experimentally confirmed miRNA-mRNA interactions inferred by each method, verified using the collection of experimentally confirmed miRNA-mRNA interactions as described in Section 2.5.

**Figure 4:**
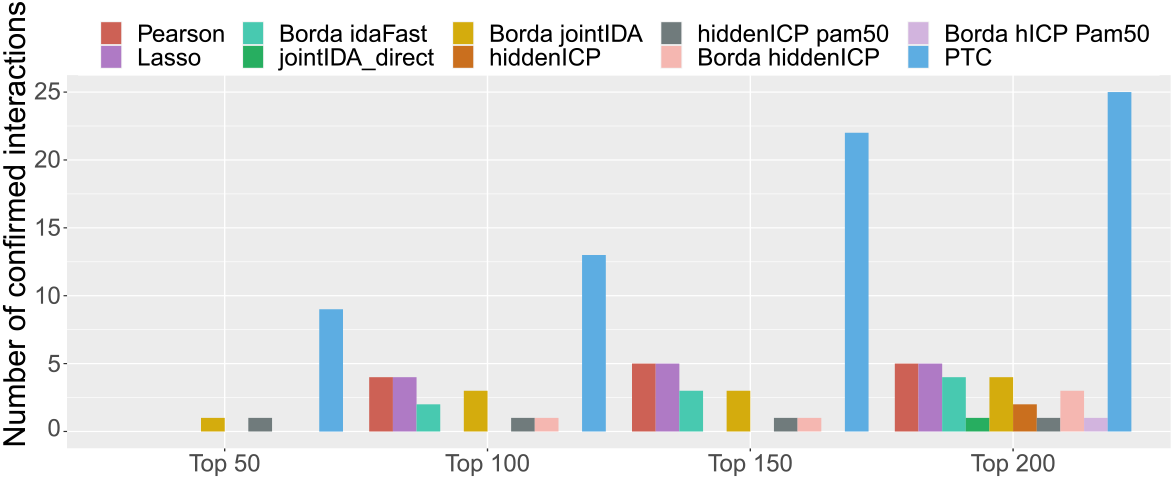
Confirmed interactions found by our method in comparison with confirmed interactions found by the other methods analyzed.

To provide the time series data required by *PTC*, the expression values of VIM in the bulk dataset were ordered and used as reference (VIM-Time) to sort the expression values of all other genes, and in this way we approximate the data set used in [18] to a pseudotime data set. Even with bulk data, which is a compendium of static data from a large number of patients rather than sequential data, *PTC* outperforms the other methods in all top lists (top 200, 150, 100 and 50). As shown in Table 2, *PTC* is able to detect 25 confirmed interactions among the top 200 (≈ 12.5%). *Pearson* and *Lasso* are the runner-up methods with 5 confirmed interactions (2.5%).

**Table 2:** Summary of performance of *PTC* using the bulk dataset, in comparison with the best two non-causal methods, the best (average) method based on static ICP and PTC. Performance was measured as the number and percentage of experimentally confirmed interactions in each top list. The number of confirmed EMT miRNA and mRNA markers inferred by each method is given in columns miR_EMT and mR_EMT respectively.

| Pearson |  |  |  |  | Lasso |  |  |  |  |
| --- | --- | --- | --- | --- | --- | --- | --- | --- | --- |
| Top | Confirmed | Percentage | miR_EMT | mR_EMT | Top | Confirmed | Percentage | miR_EMT | mR_EMT |
| 200 | 5 | 2.5% | 4 | 8 | 200 | 5 | 2.5% | 6 | 4 |
| 150 | 5 | 3.333% | 4 | 6 | 150 | 5 | 3.333% | 5 | 3 |
| 100 | 4 | 4% | 4 | 4 | 100 | 4 | 4% | 2 | 2 |
| 50 | 0 | 0% | 3 | 1 | 50 | 0 | 0% | 1 | 1 |
| Hidden ICP pam50 |  |  |  |  | PTC |  |  |  |  |
| 200 | 1 | 0.5% | 5 | 4 | 200 | 25 | 12.5% | 10 | 2 |
| 150 | 1 | 0.667% | 3 | 4 | 150 | 22 | 14.667% | 10 | 1 |
| 100 | 1 | 1% | 3 | 3 | 100 | 13 | 13% | 9 | 1 |
| 50 | 1 | 2% | 3 | 1 | 50 | 9 | 18% | 8 | 0 |

Note: Pearson and Lasso are the two methods no based in ICP with the best performance over all tops. Hidden ICP pam50 version was selected as the best (average) performance because of the fact it has confirmed predictions in all tops

Based on the hypothesis that the ranking of inferred miRNA-mRNA interactions by each method represents a measure of the strength of correlation/causal effect, the top 200, 150 100 and 50 interactions discovered by each method were analyzed, expecting an increase of the percentage of confirmed interactions. In general, the percentage of confirmed regulatory relationships increased when fewer miRNA-mRNA pairs are considered in the top. Our method was able to infer confirmed miRNA-mRNA relationship in all the top lists with a rate of at least 12.5%.

*Pearson* and *Lasso* have shown the second best performance in top 200, 150 and 100, but none of the top 50 interactions discovered by the two methods were experimentally confirmed (based on the databases used). The static version of Hidden ICP with pam50 was able to determine at least one confirmed relationship in all the top lists. In any case none of the other analyzed methods were able to obtain a percentage of confirmed interactions higher than 4%.

As an additional measurement of performance, the miRNAs and mRNAs found in the inferred regulatory relationships were analyzed to verify which of them are confirmed EMT markers. *Pearson, Lasso* and *PTC* were able to detect regulatory relationships involving 10 miRNAs that are confirmed as EMT miRNAs. Unlike the other methods, *PTC* persistently identifies relationships containing those EMT miRNAs, assigning them a high score. Because of that, relationships containing these 10 EMT miRNAs can be found in the top 200 and 150 lists of *PTC,* as shown in Table 2. The intersections of confirmed interactions inferred by all methods were also investigated. As shown in Figure 5, *PTC* was able to infer two regulatory relationships that were also inferred by at least one of the other methods. A total of 23 confirmed interactions were detected exclusively by our method.

**Figure 5:**
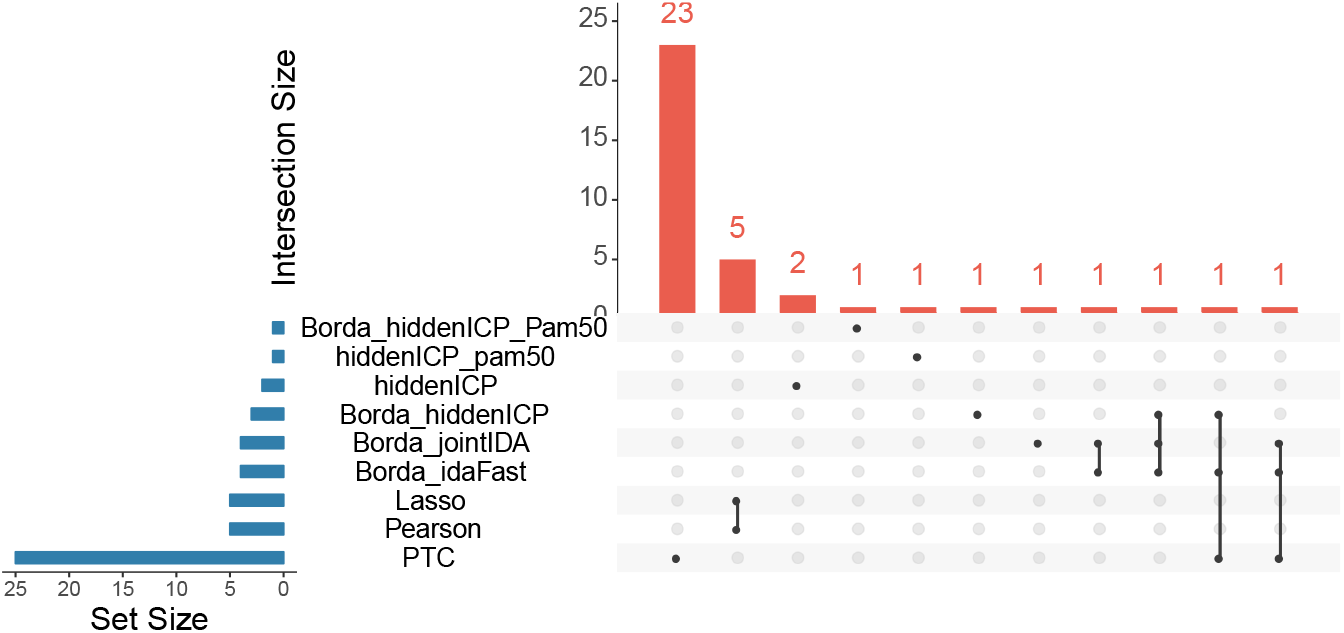
Intersection of the confirmed interactions predicted in top 200 by each method when using the bulk data [18]. The vertical axis represents the size of the intersection. Points below the horizontal axis indicate that the corresponding bar belongs to the method shown on its left. Multiple points under a bar represent the intersected methods. E.g the 25 confirmed pairs from PTC are represented as one point under the bar of 23 units long and 2 points (intersections with other methods) under the two bars of 1 unit long. Visualization made using *Intervene* [46]

## 4 Conclusion

In this paper we have presented *PTC*, a new approach for discovering causal miRNA-mRNA relationships characterising biological processes. Our method assumes that if a set of miRNAs are regulatory agents for the genes that drive a biological process, then the causal relationships between such miRNAs and mRNAs are invariant throughout the process. Based on this assumption, *PTC* performs a data transformation by means of a pseudotime analysis to create temporal ordered data, and in this way identify the invariance of the causal relationships during the biological process of interest.

In *PTC*, the pseudotime analysis phase warranties that violations to the invariance property are correctly and efficiently assessed since the test is performed on temporally ordered gene expression data. Additionally, rejecting the miRNA-mRNA pairs that are not able to biologically interact, thus they cannot have a causal relationship, help *PTC* to determine whether a miRNA-mRNA invariant relationship corresponds to a causal link throughout the process.

Experiments with the data used imply the superior performance of our method. Our method outperformed all other analyzed methods in terms of the number confirmed regulatory interactions that were predicted for each method. The results of our tests suggest that making use of the temporal information of biological processes provides additional information that significantly improves the inference of regulatory relationships. The increase in the number of confirmed relationships that were inferred by our algorithm suggests that the combination of TargetScan 7.0-based heuristics and invariance property is a powerful and effective tool for uncovering causal relationships characterising a biological process.

