## Supplementary material for "A Pseudo-Temporal Causality Approach to Identifying miRNA-mRNA Interactions During Biological Processes"

#### 1 Single cell data processing

The original single cell data set [Wang et al., 2019] was downloaded from Gene Expresson Omnibus (GEO) website, under series GSE114071. The downloaded series data contains Reads Per Kilobase Million (RPKM) values for mRNAs and log2 of the normalized reads that reflect the fraction among all miRNA reads for miRNAs. Since this dataset does not contain explicitly miRNA gene expression, we applied the transformations described below:

- **Reverse log2**: Log2-transformation was reverted by calculating 2 raised to the $miR$ power ($2^{miR}$). Where $miR$ corresponds to each value provided in the original dataset. Because of the way that data was processed by [Wang et al., 2019], the reverted values represent the fraction of a given miRNA within all miRNAs (refered as miRNA expression or miRNA levels in [Wang et al., 2019]).
- **Amplification**: Reverted data was amplified by a factor of 30.18, which is the mean of total reads that map to miRNAs for all cells reported in the original study. Duplicated miRNA names were removed by selecting each first coincidence found in GSE114071.

### 2 "GO biological processes" and "KEGG pathways" analysis

Outcome of the "GO biological processes" and "KEGG pathways" analyses for the miRNAs and genes in interactions discovered by *PTC* from the processed single cell data set [Wang et al., 2019]. These analyses were performed by using the *Enrichr* web tool [Chen et al., 2013]

Genes predicted by ***PTC*** *significantly linked with KEGG pathways and GO biological processes related with EMT and cancer*

| PTC inferred genes that are significantly associated with KEGG pathways | | | |
|---|---|---|---|
| Term | Overlap | P value | Genes |
| p53 signaling pathway | 7/72 | 0.00 | CCND3, SIVA1, GADD45A, CDK4, EI24, CDK1, CYCS |
| MicroRNAs in cancer | 13/299 | 0.00 | MIR10A, PRKCB, MIR25, MIR27B, MIR27A, MIR200C, RHOA, MIR125A, MIR423, DDIT4, MAPK1, VIM, MIR30D |
| Non-small cell lung cancer | 4/66 | 0.02 | PRKCB, GADD45A, CDK4, MAPK1 |
| Bacterial invasion of epithelial cells | 4/74 | 0.03 | ARPC2, ARPC5, RHOA, ACTG1 |
| Leukocyte transendothelial migration | 5/112 | 0.04 | RAP1B, PRKCB, MYL12A, RHOA, ACTG1 |
| **PTC inferred genes that are significantly related with GO Biological processes** | | | |
| Term | Overlap | P value | Genes |
| Regulation of hematopoietic stem cell differentiation (GO:1902036) | 10/76 | 0.00 | PSMD7, PSMA4, PSMB5, YTHDF2, PSMA1, PSMD14, UBB, UBC, H3F3A, PSMB9 |
| Regulation of hematopoietic progenitor cell differentiation (GO:1901532) | 10/77 | 0.00 | PSMD7, PSMA4, PSMB5, YTHDF2, PSMA1, PSMD14, UBB, UBC, H3F3A, PSMB9 |
| Regulation of stem cell differentiation (GO:2000736) | 10/88 | 0.00 | PSMD7, PSMA4, PSMB5, YTHDF2, PSMA1, PSMD14, UBB, UBC, H3F3A, PSMB9 |
| Negative regulation of cell migration involved in sprouting angiogenesis (GO:0090051) | 2/14 | 0.02 | PDCD10, RHOA |
| Regulation of cell division (GO:0051302) | 4/65 | 0.02 | PRC1, TXNIP, PIN1, CALM2 |
| Regulation of cell differentiation (GO:0045595) | 5/121 | 0.05 | DDX5, PPP2R1A, PRKCB, PA2G4, ANP32B |

A large number of genes predicted by or method are significally related with enriched terms of EMT processes. Note: P value = 0.00 means P value $< 0.01$, Overlap column is defined as the number of inferred genes that are in the term/all genes of that term
